# NLRP7 Plays A Functional Role in Regulating BMP4 Signaling During Differentiation of Patient-Derived Trophoblasts

**DOI:** 10.1101/850420

**Authors:** Aybuke Alici-Garipcan, Burcu Özçimen, Ilke Suder, Volkan Ülker, Tamer T. Önder, Nesrin Özören

## Abstract

Complete Hydatidiform Mole (HM) is a gestational trophoblastic disease resulting in hyper proliferation of trophoblast cells and absence of embryo development. Mutations in the primate specific-maternal effect gene NLRP7 are the major cause of familial recurrent complete HM. Here, we established an in vitro model of HM using NLRP7 deficient patient-specific induced pluripotent stem cells (iPSCs) derived trophoblasts. Using whole transcriptome profiling during trophoblast differentiation, we showed that NLRP7 deficiency results in precocious downregulation of pluripotency factors, activation of trophoblast lineage markers and promotes maturation of differentiated extraembryonic cell types such as syncytiotrophoblasts. Interestingly, we found that these phenotypes are dependent on BMP4 signaling and BMP pathway inhibition corrected the excessive trophoblast differentiation of patient derived iPSCs. Our human iPSC model of a genetic placental disease recapitulates aspects of trophoblast biology, highlights the broad utility of iPSC-derived trophoblasts for modeling human placental diseases and identifies NLRP7 as an essential modulator of key developmental cell fate regulators.

## Introduction

Maternal effect genes encode for proteins deposited into oocytes that are required for embryonic development (1). Mutations in such genes result in embryonic phenotypes that reflect the genotype of the mother rather than that of the offspring (2). In mice models, null phenotypes of a majority of these genes result in arrested development at very early embryonic time points (3). Familial biparental hydatidiform mole (FBHM; MIM 231090) is the only known pure maternal-effect inherited disorder in humans. Complete molar pregnancy (CHM) is a gestational trophoblastic disease, defined as a pregnancy with no embryo along with abnormal hyper-proliferation of extraembryonic trophoblastic tissue (4, 5). The major cause for FBHM is homozygous or compound heterozygous maternal-effect mutations in NLRP7 (NOD-like receptor family pyrin domain containing 7) (6–9). Certain NLRP7 mutations found in a heterozygous state have also been associated with recurrent reproductive failure (10). Interestingly, two of the first mammalian maternal effect genes identified encode for NLR proteins, Nlrp5 (Mater) and Nlrp2 both of which are required for early embryonic development of mice and have unknown functions (11, 12). In contrast to mice models of maternal effect genes, embryos of affected women with NLRP7 mutations do not arrest at a very early stage, but rather undergo excessive differentiation and commitment to extraembryonic lineages in vivo. The NLRP7 gene is present only in primates (13, 14). NLRP7 has been reported to be involved in inflammasome formation and cytokine release (15, 16) as well as in modulating DNA methylation and trophoblast differentiation (17, 18). FBHM associated NLRP7 mutations have been linked to imprinting abnormalities (19). However, how NLRP7 regulates early embryonic development and how its absence contributes to human disease pathogenesis remain unclear. Due to its primate-specific nature, it has not been possible to study the role of this maternal effect gene during embryogenesis in model organisms. Patient-derived disease specific induced pluripotent stem cells (iPSCs) provide a powerful platform for overcoming the ethical and technical challenges associated with studying early human embryogenesis. To derive trophoblasts from hESCs several groups have reported the feasibility of BMP4 treatment, alone or in combination with inhibitors of FGF and ACTIVIN/NODAL signaling (20–22). Here, we established iPSCs-derived trophoblasts from an HM patient with NLRP7 deficiency and showed that these cells provide a model to investigate human placental diseases. Finally, our work demonstrates that NLRP7 plays a pivotal role in trophoblast differentiation by modulating the BMP4 pathway.

## Results

### Generation of Human Patient derived iPSC from HM patient with NLRP7 deficiency

To assess the function of NLRP7 in HM pathology, we generated iPSC from a patient carrying compound heterozygous variant of NLRP7 with a prior diagnosis of recurrent HM (23, 24) (Figure 1A). Fibroblasts form a HM patient and an unrelated healthy control (WT) were reprogrammed using non-integrating episomal plasmids(25, 26). No difference was observed between HM and WT cells in terms of reprogramming efficiency and maintenance of pluripotency under standard culture conditions (Figure 1B-D). iPSCs expressed pluripotency markers OCT4, NANOG, SOX2 LIN28A and LIN28B as assessed by immunofluorescence and RT-PCR (Figure 1C and 1D). iPSC lines were karyotypically normal, devoid of episomal reprogramming vectors, and could readily differentiate into cell types belonging to three germ layers in a teratoma formation assay in immunocompromised mice (Figure S1A-C). We confirmed NLRP7 deficiency in HM cells at the molecular level (Figure 1E, 1F, and S1D). We observed that reprogramming promoted the expression of NLRP7 in WT cells, while HM^iPSC^ had significantly reduced expression (Figure S1E). On the other hand, NLRP2, the closest homologue of NLRP7, levels were not affected in patient-derived iPSCs (Figure S1D and 1E). These results indicate that somatic NLRP7 mutant cells can be successfully reprogrammed to pluripotency via exogenous transcription factors.

**Fig. 1.**
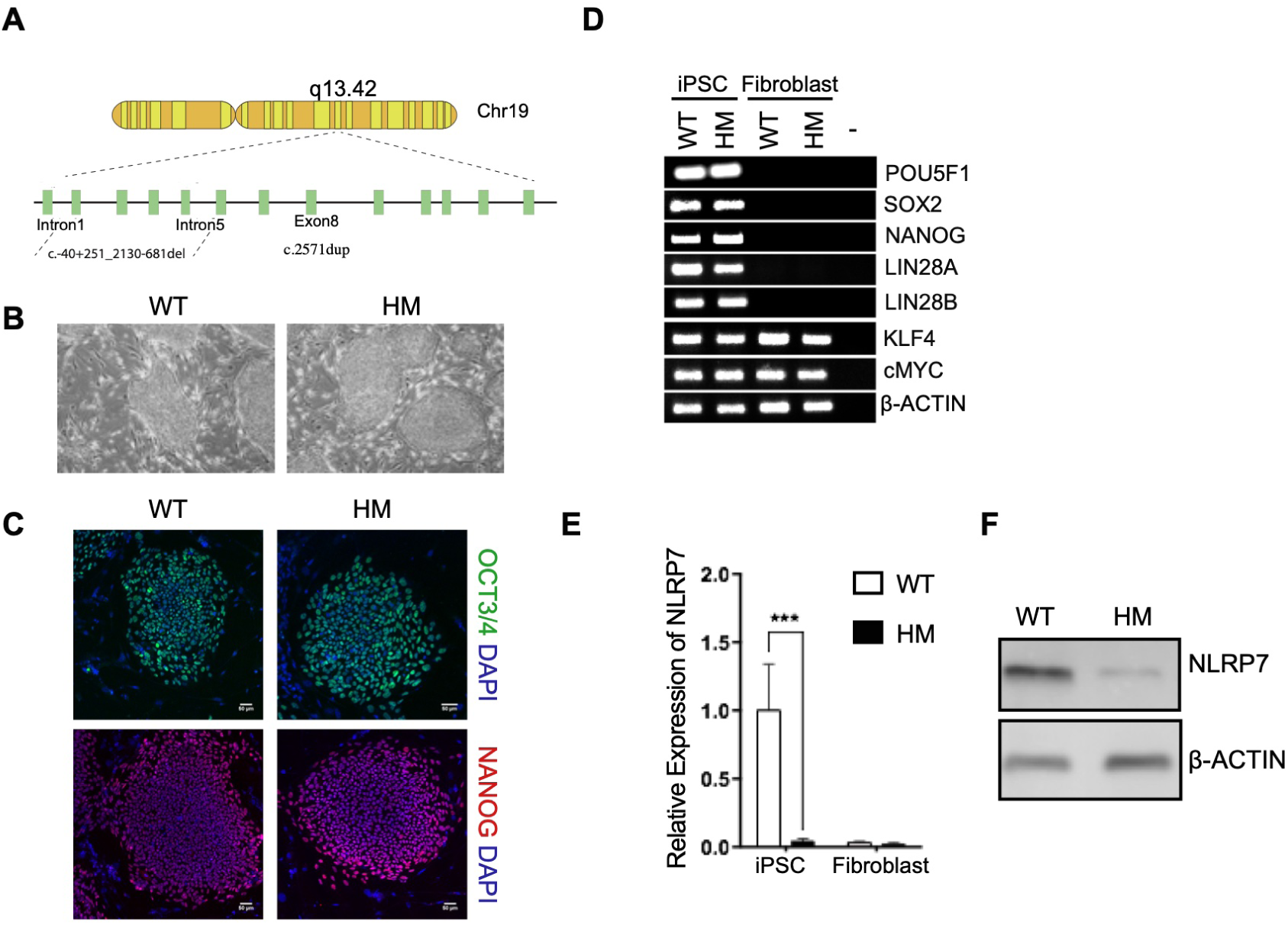
Characterization of HM-specific iPSCs carrying NLRP7 mutations. (A) Schematic and coordinates of the deletion and the single base pair duplication on NLRP7 gene in HM cells used in this study. (B) Colony morphologies of reprogrammed cells on MEFs under light microscopy. Images were acquired at 4X magnification. (C) Immunostaining for pluripotency markers OCT3/4 and NANOG. DAPI was used as a nuclear stain. (D) RT-PCR showing the expression of pluripotency markers in WT and HM iPSCs and their fibroblast counterparts. (E) NLRP7 mRNA levels in WT and HM iPSCs and their fibroblast counterparts. Relative mRNA levels were normalized to GAPDH. n = 3 biological replicates. The bars represent mean ± SD. \*\*\**P* < 0.005 by 2-way ANOVA followed by Sidak’s multiple comparison test. (F) NLRP7 protein levels in WT and HM iPSCs as detected by immunoblotting. *β*-actin was used as a loading control.

### NLRP7 Deficient iPSC Derived Trophoblasts Closely Resemble HM Pathogenesis

Next, we generated trophoblast cells from iPSCs using an established protocol of BMP4 treatment in combination with inhibition of ALK5 and FGF signaling (via A83-01 and PD173074; referred to as BAP condition) (22, 27, 28) (Figure 2A). On the second day of BAP exposure, WT cells became distinguishable by their flattened morphology from undifferentiated counterparts. Intriguingly, HM cells started to lose their distinct iPSC morphology only after 24 hours of BAP treatment (Figure 2B). To characterize the differentiation of iPSCs towards trophoblast-like cells at the whole transcriptome level, we carried out RNA-sequencing analysis of WT^iPSC^ and HM^iPSC^ over the time course of differentiation. Hierarchical clustering analysis pointed to a high degree similarity between the WT^iPSC^ and HM^iPSC^ indicating that NLRP7 mutations do not lead to significant transcriptional changes in the pluripotent state (Figure S2A). Upon BAP treatment, the transcriptomes of the groups differed from their parental iPSCs. To address the extent of trophoblast differentiation in our culture system, we compiled a list of general trophoblast markers based on the recent literature (Table S1). Gradual up-regulation of trophoblast genes was observed during the time course of differentiation of WT cells upon BAP exposure indicating the success of BAP to direct commitment of pluripotent stem cells to trophoblast lineage (Figure S2B). Gene Set Enrichment Analysis (GSEA) further confirmed highly significant downregulation of ESC-specific genes and upregulation of placental genes (Figure S2C). At the whole transcriptome level HM^BAP^ cells were enriched in trophoblast signature whereas pluripotency signatures were attenuated in comparison to WT^BAP^ cells in a time dependent manner (Figure 2C-E, S2E). Of note, the transcripts of many trophoblast cell fate transcription factors were upregulated in HM^BAP^ such as TFAP2C, GATA3, CDX2 (29, 30). Although no difference was observed in reprogramming efficiency of iPSC from WT and HM cells, the transcripts of key transcription factors regulating trophoblast lineage commitment; EOMES, GATA4 and ITGA4 were remarkably elevated in HM^iPSC^ under self-renewal conditions. To further assess the differentiation efficiency, we analyzed widely used trophoblast markers: CDX2, HLA-G, KRT7 and the pluripotency marker, OCT3/4 protein expression by immunostaining. Although, BAP-treated cells of both groups started to be positive for early trophoblast lineage marker, CDX2 on day 2, HM cells tended to gain more CDX2 positivity than WT cells (Figure 2F). On the other hand, HLA-G positive cells were visible mostly where CDX2 was absent on day 4 indicating the later stages of differentiation. Staining for HLA-G was considerably stronger and more uniform in HM cells. Although, the majority of the cells from both groups were positive for KRT7 even on day 2, HM cells stained more strongly than WT cells for all time points assessed. Consistent with the initiation of differentiation, OCT3/4 staining declined after BAP exposure for 2 days, with HM cells exhibiting a greater degree of OCT3/4 loss than WT cells. We also noticed that HM cells became enlarged and more uniform when compared to WT cells in response to BAP treatment. As, increased cell size is an initial event in trophoblastic differentiation (31, 32), we measured the sizes of DAPI stained-cell nuclei. HM cells had significantly larger nuclei than that of WT cells (Figure S2E). In accordance with immunostaining results, OCT3/4 was not detectable on day 4 of BAP treatment, whereas CDX2, KRT7 and HLA-G were elevated in HM cells upon BAP exposure (Figure 2G). Placental growth factor (PGF) is a placental hormone produced predominantly by functional trophoblasts during pregnancy (33). HM cells produced significantly more soluble PGF than WT cells on day 4 (Figure 2H). We then tested whether BAP treatment resulted in the differentiation of a specific sub-type of trophoblasts, namely, cytotrophoblasts (CT), extravillous trophoblasts (EVT) and syncytiotrophoblasts (ST). We compiled lists of markers indicative of different trophoblast populations (Table S2) and examined their expressions during differentiation. While CT and EVT marker genes were almost similarly induced in WT^BAP^ and HM^BAP^ cells, ST related genes such as ERVW-1 (Syncytin) were remarkably enriched in HM^BAP^ cells. As STs are terminally differentiated trophoblasts, HM cells appeared to exhibit a more advanced differentiation status (Figure S2F). Taking together, these results suggest that patient-specific iPSC derived trophoblasts can successfully recapitulate HM disease pathogenesis and highlight an early activation of trophoblast lineage commitment dependent on NLRP7 deficiency.

**Fig. 2.**
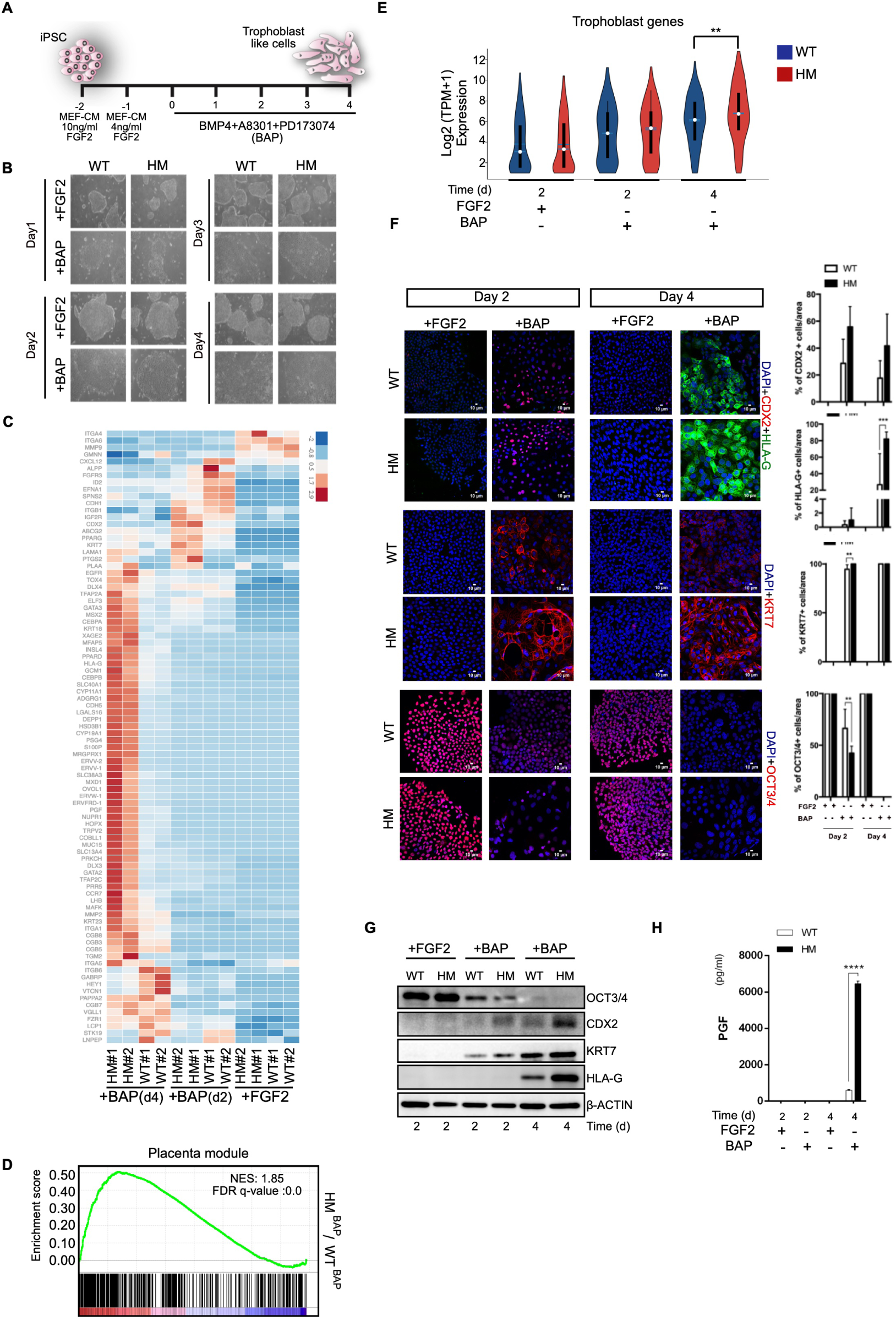
NLRP7 deficiency promotes trophoblast differentiation. (A) Diagram of the trophoblast differentiation protocol. (B) Changes in colony morphologies upon BAP exposure. Images were acquired at 4x magnification. (C) Heatmap showing the expression of trophoblast markers across the time course of differentiation of WT and HM iPSCs (n=2 biological replicates) (D) Gene set enrichment analysis (GSEA) of Placenta module. Genes were ranked according to log2 fold changes in gene expressions comparing HM^BAP^ cells to WTBAP cells on day 4. (E) Violin plot displaying log-transformed expression of trophoblast genes in 2C (n=2, p <0.05, Wilcoxon rank sum test). (F-G) Immunostaining and immunobloting for trophoblast markers, CDX2, HLA-G, KRT7 and stem cell marker, OCT3/4. (F) Percentage of the cells immunostained for CDX2, HLA-G, KRT7 and OCT3/4. Scale bars; 10µM. (G) Representative immunoblots for CDX2, HLA-G, KRT7 and OCT3/4. (H) 24-hours PGF production as assessed by ELISA. (F and H) ±SD, n = 3, *p ≤ 0.05, ***p < 0.005, ****p < 0.001 by 2way ANOVA followed by Sidak’s multiple comparison test.

### BMP4 Independent Derivation of Trophoblasts from NLRP7 Deficient iPSCs

As BMP4 is known to induce trophoblast differentiation, we investigated whether BMP4 signaling could be responsible for the phenotype observed in HM cells. We removed BMP4 from the differentiation medium (AP) and assessed the predisposition of HM cells to differentiate to trophoblasts. Surprisingly, HM^AP^ cells displayed predominantly homogeneous phenotype resembling the morphology of BAP treated cells on day 4 (Figure S3A). WT cells did not exhibit such a difference in morphology from undifferentiated iPSCs. Hierarchical clustering of whole transcriptome data demonstrated that HM^AP^ cells were broadly distinct from WT^AP^ cells on day 4 (Figure S3B). Remarkably, HM cells expressed genes related to trophoblasts to a substantial level when treated only with AP in the absence of BMP4 (Figure 3A) and the differentially expressed transcripts in HM^AP^ cells were highly associated with the placenta (Figure 3B). Moreover, the global expression of trophoblast genes was enriched in HM^AP^ cells (Figure 3C), whereas the transcripts of pluripotency genes declined in comparison to WT^AP^ cells (Figure S3C). We verified that transcripts of CDX2 on day 2 and PGF, INSL4, PSG on day 4 were higher in HM cells compared to WT cells (Figure 3D). On the other hand, no significant global upregulation was observed for the genes related to endoderm, mesoderm or mesoendoderm differentiation (Figure S3D). Immunostaining revealed that CDX2 positive cells were evident in HM group on day 2 and HLA-G positive cells emerged on day 4 upon AP treatment (Figure 3E). These results mirrored the phenotype observed for that of BAP exposure albeit at a lower level for the trophoblast specific markers. All of the HM cells strongly stained for KRT7 on day 4 unlike WT^AP^ cells. HM cells also exhibited bigger nuclei than WT cells upon AP exposure (Figure S3E). Western blot analysis confirmed that AP treated HM cells expressed substantially more CDX2, KRT7 and HLA-G on day 4 compared to WT cells (Figure 3F). OCT3/4 protein levels decreased dramatically on day 4 in HM cells showing that HM cells lost pluripotency properties faster than WT cells. PGF secretion provided a clear evidence for the trophoblastic features of HM^AP^ cells (Figure 3G). We next examined whether NLRP7 deficiency could endow HM cells with the capacity to differentiate into terminal sub-types of trophoblasts under AP conditions. HM^AP^ cell population contained CT, EVT and ST-like cells as evidenced by the robust expression of trophoblast markers belonging to these subtypes (Figure S3F). Trophoblast differentiation in NLRP7 deficient HM cells in the absence of BMP4 suggested that HM cells may undergo excessive trophoblast differentiation due to dysregulation of BMP4 signaling.

**Fig. 3.**
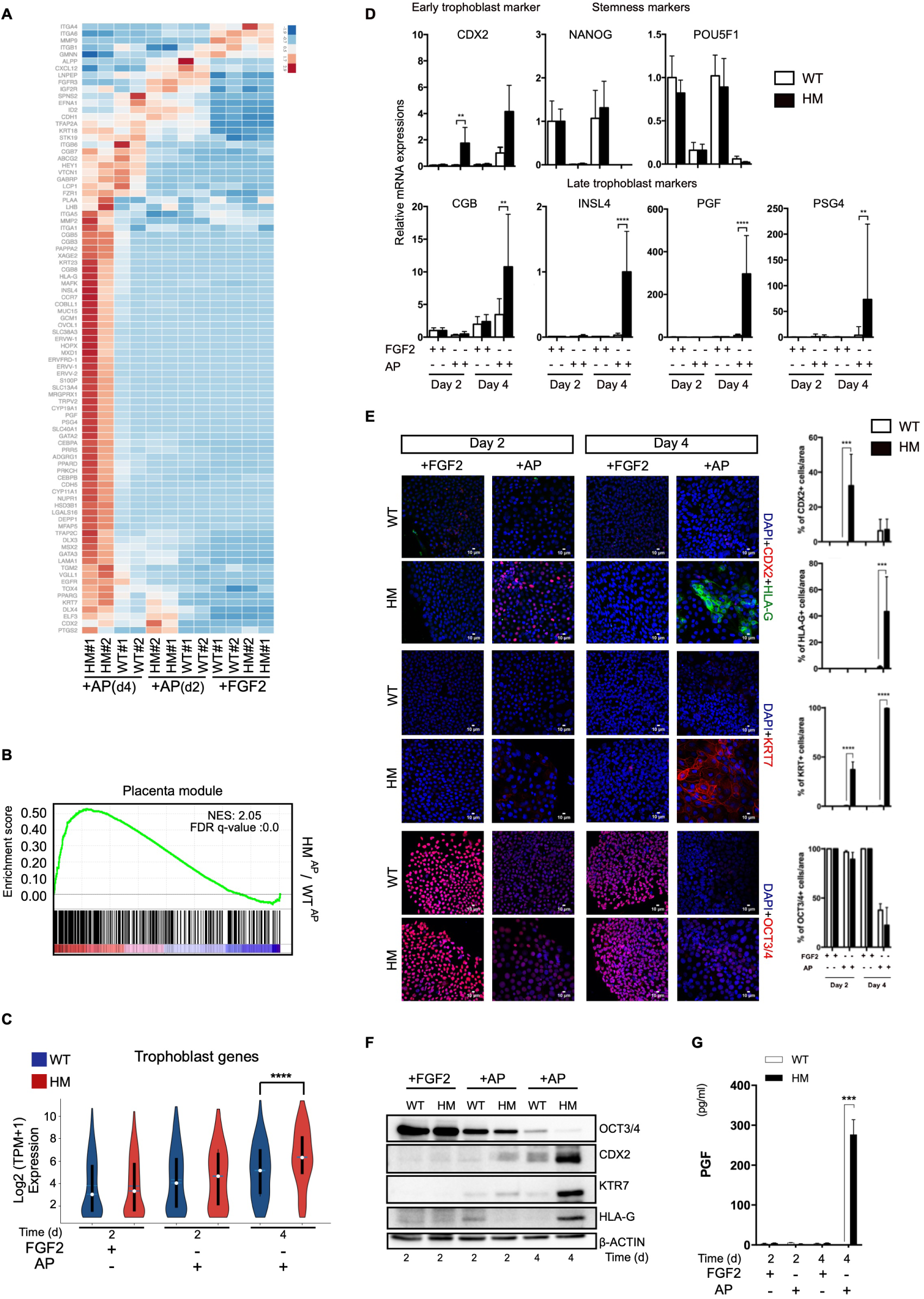
NLRP7 deficiency obviates the requirement for exogenous BMP4 during trophoblast differentiation. Cells were treated with either FGF2 as a control or AP (A) Heatmap depicting the expression of trophoblast markers during time course of WT and HM iPSC differentiation (n=2 biological replicates). (B) GSEA of Placenta module. Genes were ranked according to log2 fold changes in expression comparing HM^AP^ cells to WT^AP^ cells on day 4. (C) Violin plot displaying log-transformed expression levels of trophoblast genes in Figure 3A (n=2, p value <0.05, Wilcoxon rank sum test). (D) RT-qPCR for trophoblast and stem cell markers. (E and F) Immunostaining and immunoblotting for CDX2, HLA-G, KRT7 and OCT3/4. (E) Percentage of the cells immunostained for CDX2, HLA-G, KRT7 and OCT3/4. Scale bars; 10µM. (F) Representative immunoblots for CDX2, HLA-G, KRT7 and OCT3/4. G) 24-hours PGF production as assessed by ELISA. (D,E and G) *p ≤ 0.05, ***p < 0.005, ****p < 0.001 by 2way ANOVA followed by Sidak’s multiple comparison test. The bars represent mean ± SD, n = 3 biological replicates.

### NLRP7 Modulates BMP4 Signaling during Trophoblast Differentiation

Promotion of trophoblast lineage commitment without exogenous BMP4 led us to hypothesize that NLRP7 may act on trophoblast differentiation through BMP-4 signaling. Interestingly, RNA-sequencing revealed that BMP signaling related genes were significantly enriched in the transcriptome of HM cells when treated with AP at the early time point (Figure 4A). We uncovered that BMP4 mRNA levels and soluble BMP4 secretion were higher in HM^AP^ cultures in comparison to WT^AP^ cultures (Figure 4B and 4C). As, aberrant DNA methylations have been observed in recurrent familial moles (34), we investigated whether increased BMP4 expression of HM cells is stemmed from altered methylation status. However, we could not find a correlation between methylation status and expression levels of BMP4 (Figure S4). To investigate active BMP4 signaling, we generated a list of BMP4-responsive genes using our whole transcriptome data of BAP-treatment in comparison to AP-treatment of WT cells on day 2 (200 number of genes, Table S3). In HM^AP^ cells, we observed that upregulation of this set of BMP4-responsive genes independent of BMP4 exposure (Figure 4D). Moreover, expression of BMP4 targets, GATA-2 and GATA-3, which are also important transcription factors for trophoblast lineage commitment during embryogenesis, was significantly higher in HM cells (Figure 4E). Western blot analysis confirmed markedly increased phosphorylation of pSMAD1/5/9 in HM^AP^ cells during the time course of differentiation indicating an overactive BMP signaling as the underlying mechanism of augmented trophoblast differentiation of HM^AP^ cells (Figure 4F). Taken together, these results point to a reciprocal association between NLRP7 and BMP4 during trophoblast differentiation and suggest that hyper-trophoblast differentiation observed in HM cells with NLRP7 deficiency is due to aberrant BMP4 signaling.

**Fig. 4.**
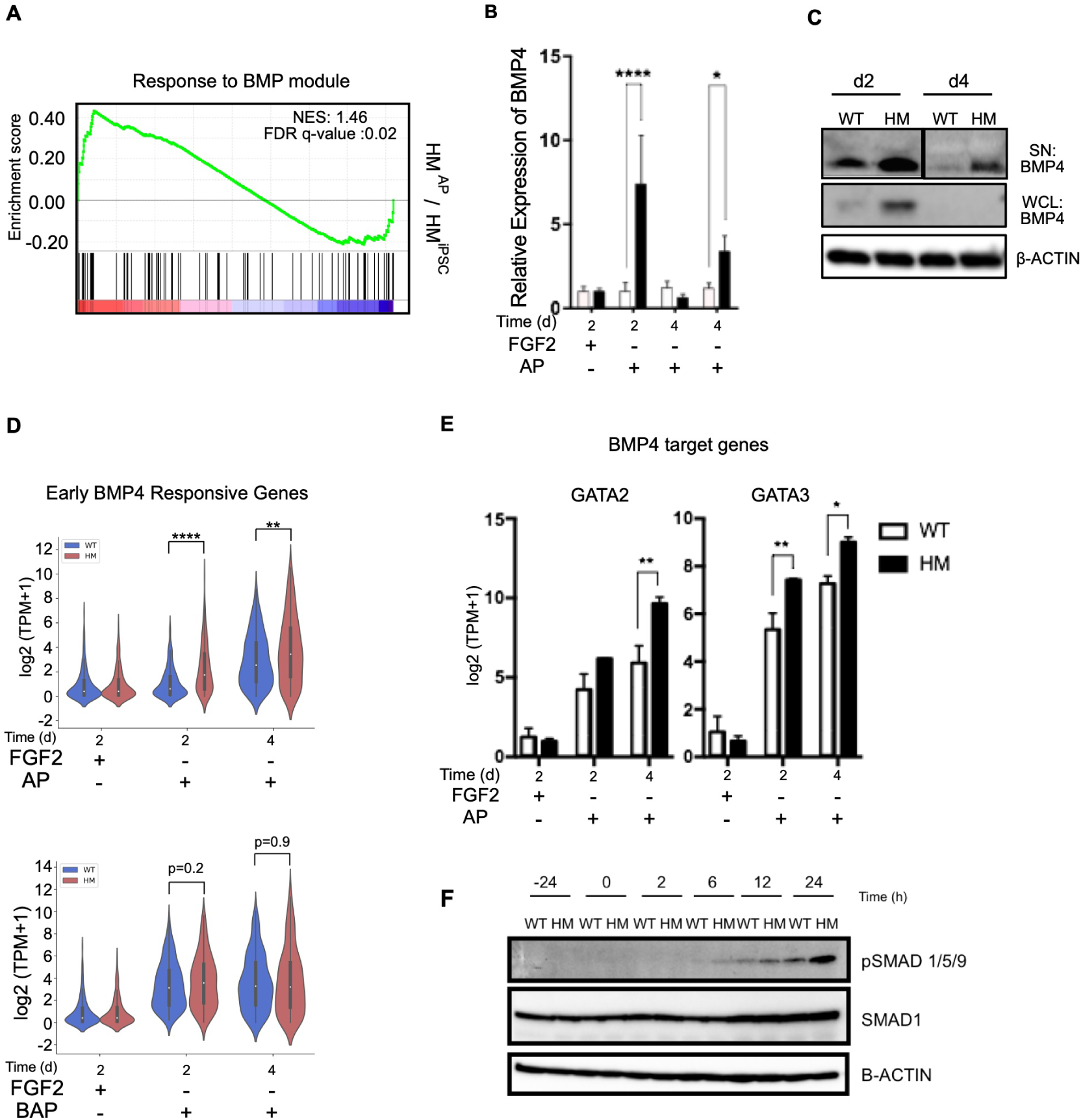
NLRP7 deficiency modulates BMP signaling under AP conditions. (A) GSEA showing the enrichment of Response to BMP gene set in HM^AP^ cells in comparison to HM^iPSC^. Genes were ranked according to log2 fold changes in gene expressions comparing HM^AP^ cells to HM^iPSC^ cells on day 2. (B) RT-qPCR showing the expression of BMP4. n = 3 biological replicates. (C) Western blotting of BMP4. Supernatant, SN; whole cell lysate, WCL. (D) Violin plots for early BMP4 responsive genes in WT and HM cells across the time-course of differentiation (p<0.05, Wilcoxon rank sum test). (E) Log-transformed expression of BMP4 target genes, GATA2 and GATA3. (A,D,E) n=2 biological replicates. The bars represent mean ± SD. (B,E) *p ≤ 0.05, **p < 0.01, ***p < 0.005, ****p < 0.001 by 2way ANOVA followed by Sidak’s multiple comparison test. F) Western blotting of pSMAD1/5/9 and total SMAD1.

### NLRP7 Reintroduction or Inhibition of BMP Pathway Revert the Excessive Trophoblast Differentiation of Patient Specific iPSCs

Next, we evaluated whether correction of NLRP7 deficiency or suppression of BMP4 signaling could restore abnormal trophoblast differentiation of HM cells. To test this, NLRP7 was stably over-expressed in HM cells (HM ^+NLRP7^) (Figure S5A). Overexpression of NLRP7 not only led the downregulation of BMP4 and CDX2 proteins, but also restored OCT3/4 levels, indicating rescue of HM differentiation phenotypes (Figure S5B and S5C). Next, we evaluated if an inhibitor of BMP receptors ALK2 and ALK3 (LDN193189) would be able to attenuate trophoblast differentiation. LDN193189 exposed HM cells showed a similar morphological phenotype to WT cells (Figure 5A). At the whole transcriptome level LDN193189 treated cells clustered with undifferentiated iPSCs rather than AP treated cells (Figure S5D). Heatmap for trophoblast markers also clearly demonstrated that BMP pathway inhibition impeded the differentiation of HM cells (Figure 5B). The global expression of trophoblast markers was also reduced in response to LDN193189, whereas pluripotency markers remained highly expressed (Figure 5C and 5D). We further confirmed the dramatic decline in trophoblast gene expression upon BMP pathway inhibition by qRT-PCR (Figure 5E). In concordance with transcriptomic analyses, BMP pathway inhibition blocked the upregulation of CDX2, KRT7, HLA-G and prevented the downregulation of OCT3/4 in HM cells (Figure 5F). Overall, these results indicate that in HM cells, exit from pluripotency and rapid differentiation tendency to trophoblasts is associated with aberrant BMP signaling.

**Fig. 5.**
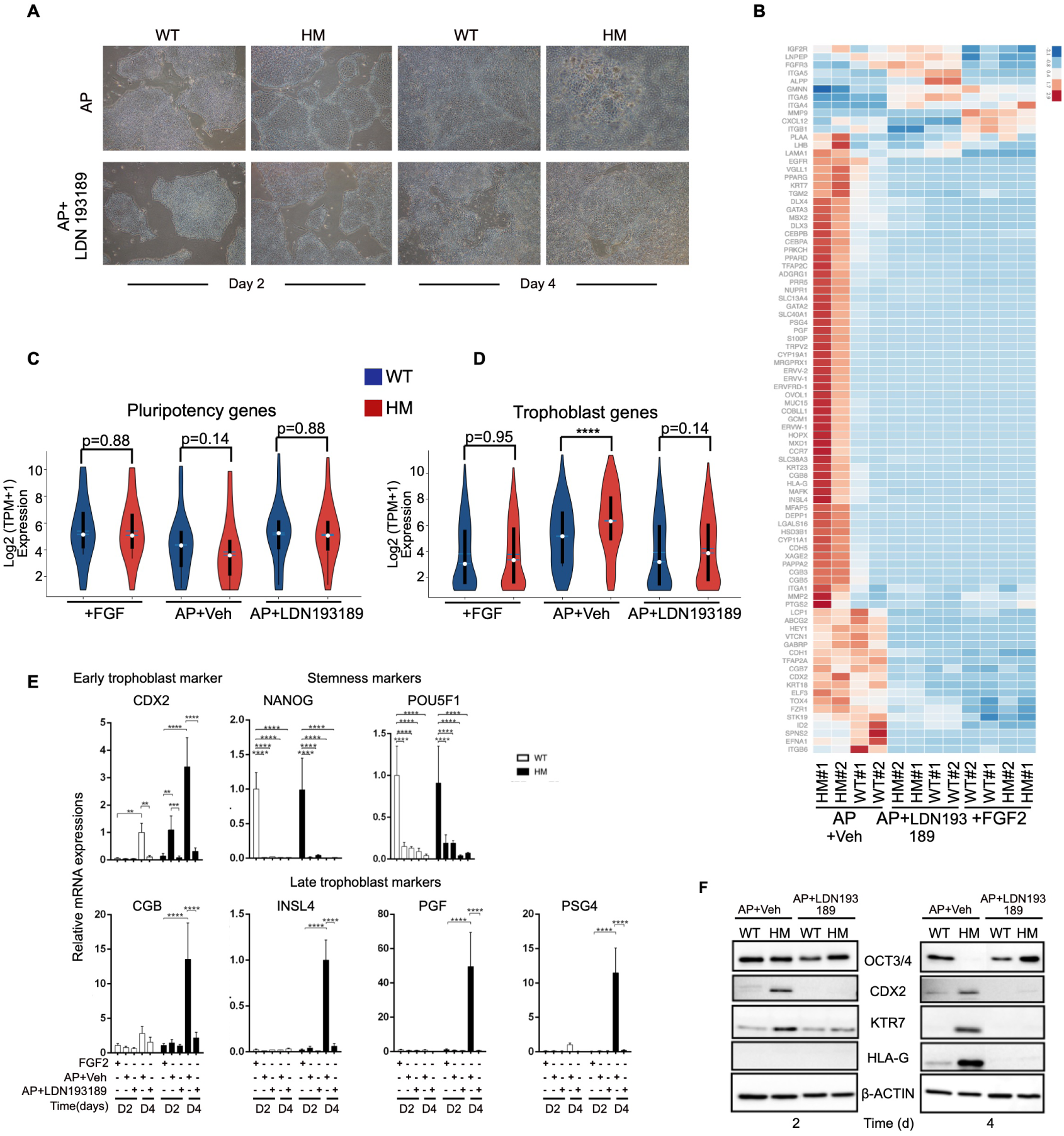
Rescue of elevated trophoblast differentiation in HMiPSC by inhibition of BMP signaling. Cells were exposed to vehicle (DMSO) or BMP signaling inhibitor LDN193189 (100nM) under AP conditions. (A) Changes in colony morphologies upon LDN193189exposure. Images were acquired at 4x magnification. (B) Heatmap showing the expression of trophoblast markers across the time-course of differentiation of HM and WT iPSCs. (C, D) Violin plots for pluripotency (C) and trophoblast (D) markers. (n=2, **** p <0.001, Wilcoxon rank sum test). (E) RT-qPCR for trophoblast and stem cell markers. *p ≤ 0.05 **p < 0.01 ***p < 0.005 ****p < 0.001 by 2way ANOVA followed by Sidak’s multiple comparison test. (F) Representative immunoblots for CDX2, HLA-G, KRT7 and OCT3/4.

## Discussion

Here, we established a patient-specific iPSC-based HM model by generating trophoblast-like cells from iPSCs carrying NLRP7 deficiency via BMP4 treatment and inhibition of FGF and Activin/NODAL signaling pathways. Both WT and HM-derived trophoblast cells were able to represent characteristics of trophoblast lineage consistent with previous reports (20, 22, 27, 28, 35, 36). The association of NLRP7 with trophoblast differentiation has been previously reported in H9 human ESCs (17) and primary trophoblasts (18). Similarly, we showed expedited trophoblast differentiation in NLRP7 deficient-patient derived HM cells upon BAP exposure compared to WT cells. Similar to hyperplasia of trophoblast tissue in Hydatidiform moles, HM^iPSC^ cells exhibited augmented trophoblast lineage commitment upon BAP exposure. Thus, our model recapitulates salient features of this gestational trophoblastic disease. We investigated the molecular mechanisms that drive cells to the trophoblast fate and uncovered an association between NLRP7 deficiency and excessive BMP4 signaling. We found that HM^iPSC^ were still able to differentiate to trophoblasts even in the absence of BMP4, due to endogenously elevated BMP4 signaling in HM cells. Suppression of excessive trophoblast differentiation by the BMP pathway inhibitor, LDN193189, verified the involvement of BMP pathway in trophoblast lineage fate driven by NLRP7 deficiency in HM cells. Notably, NLRP7 regulates the expression of key transcription factors functioning in first cell fate decision during early embryogenesis such as; GATA3 and CDX2. NLRP7 has been shown to interact with the transcription factor YY1 (17). YY1 can repress BMP signaling (37, 38) and BMP4 is known to induce CDX2 expression (39). Hence, it is tempting to speculate that NLRP7 may restrain BMP4 levels through its interaction with YY1. As no embryo formation is observed in FRCHM, the deficiency of NLRP7 may impair ICM formation through elevated BMP4 levels by facilitating trophoblast lineage commitment. Thereby, all of the cells during the first cell fate decision may be directed to the trophoblast fate. In agreement with this, recent studies with *in vitro* pre-implantation embryos have verified the requirement of NLRP7 for proper embryo development(40, 41). Interestingly, NLRP7 was found to be one of the most upregulated genes in embryonic carcinomas, epiblasts and naive pluripotent stem cells along with core stem cell markers, such as; OCT3/4, NANOG (42–44) in correlation with our results that reprogramming process itself boosts NLRP7 expression and NLRP7 deficient cells tend to lose pluripotency properties when the media is not supplemented with FGF2. While successful pregnancy requires maternal NLRP7, its deficiency is not incompatible with pluripotency under standard ESC conditions, but once the cells exit from it, differentiation to trophoblast lineage is heavily favored. Taken together, our results show that NLRP7 deficient iPSCs can faithfully model complete Hydatidiform moles. NLRP7 deficiency predisposes pluripotent cells to the trophoblast lineage commitment by regulating the BMP4 pathway. Importantly, BMP pathway inhibition ameliorates trophoblast differentiation due to NLRP7 deficiency, which may pave the way to therapeutic treatments for HM patients.

## Materials and Methods

### Primary culture from human skin biopsy

The skin samples were obtained from a 30 year old female patient and a 44 year old healthy volunteer with informed consent under approval by Bogazici University Human Research Institutional Review Board (2014/34) by a 3 mm punch biopsy after injection of local anesthetic. The skin samples were transferred in DMEM (GIBCO) on ice. Then, the samples were washed with PBS. After washing step, skin samples were cut into relatively small pieces by a surgical scalpel. The smaller skin samples were placed on 6 well plate. Then, 22mm glass cover slips were placed onto samples to stabilize them in 4oC complete DMEM (DMEM supplemented with 10% FBS (GIBCO), 2 mM L-Glutamine (GIBCO), 1X MEM Non-Essential Amino acids (GIBCO), 100 U/ml penicillin and 100 µg/ml (GIBCO) and incubated at 37oC and 5% CO2 incubator. Medium was changed every 3-4 days. 10% DMSO 10% FBS 80% complete DMEM was used to freeze the cells.

### Patient specific iPSC generation

Patient specific iPSCs were established by episomal transfection previously described elsewhere (25, 45). The day before the episomal transfection of reprogramming vectors, primary fibroblast cells were seeded into 6 well plates at a density of 3×10^5^ and incubated overnight at 37 °C, 5% CO_2_. 1µg of each plasmid pCXLE-Oct3/4-shp53, pCXLE-SK, pCXLE-UL, pCXWB-EBNA were transfected and CXLE-eGFP, pCXWB-EBNA were used as a transfection control by NeonQR Transfection System (Thermo Scientific) via electroporation with 1400V, 20ms, 2pulse. Six days later from transfection, mitomycin-c (Sigma) treated MEFs were seeded on 6-well plates coated with 0.2% gelatin (Sigma). On the next day (day 7), reprogrammed cells were harvested and transferred to plates containing MEFs. Then, the medium was changed with hES medium containing 10 ng/mL of FGF2 (Peprotech) every other day.

### EBNA integration assay

Genomic DNA from iPSCs was isolated using a commercial kit (MACHEREY-NAGEL) as indicated by the manufacturer. PCR was performed using 50 ng genomic DNA as template per reaction with the following primers: EBNA-Fwd: AGGGCCAAGACATAGAGATG, EBNA-Rev: GCCAATGCAACTTGGACGTT, GAPDH-Fwd: ATCACCATCTTCCAGGAGCGA, GAPDH-Rev: TTCTCCATGGTGGTGAAGACG. PCR products were sequenced by Macrogen Inc. (Korea).

### Teratoma formation assay

Three confluent 10 cm plate were detached using regular passaging protocols and resuspended in 50% Matrigel (Corning) and 50% cold DMEM medium supplemented with 10% FBS, 2 mM L-Glutamine, 100 U/ml penicillin and 100 µg/ml streptomycin and keep on ice. The mixture was applied to tree SCID mice via intramuscular injection under anesthesia. The anesthetized mice were sacrificed via an IACUC-approved method after 6-8 weeks of injection and teratoma was dissected out. Teratoma was fixed in 10% formalin. Histopathological staining and examination were performed.

### Trophoblast differentiation

iPS cells were maintained routinely on MEFs with hES media (10 ng/mL FGF2). For the trophoblast differentiation, 2.4 × 10^4^ cells per square centimeter were seeded on matrigel coated plates with conditioned hES medium by a monolayer of mitomycin-C treated MEF feeder cells (MEF-CM) containing FGF2 (10 ng/mL). On the next day, medium was changed to MEF-CM containing 4 ng/mL of FGF2. Next day, the medium was changed to BMP4 (10 ng/mL) (RD Systems), the ALK4/5/7 inhibitor, A83-01 (1µM) (Tocris), and the FGF2-signaling inhibitor PD173074 (0.1 µM) (Sigma) containing (BAP) hESC basal medium not conditioned with MEF feeder cells (22, 27, 28). Control cultures were grown in the presence of FGF2 and in the absence of BAP. The medium was replenished daily.

### Quantitative Real Time-PCR

Total RNA was extracted by Direct-zol RNA Isolation Kit (Zymogen) and cDNA was synthesized by Sensifast cDNA synthesis kit (Bioline) as described by the manufacturer. Primers were synthesized by Macrogen as listed in Supplementary table 4. qRT-PCR was performed with SensiFASTTM SYBRQR No-ROX Kit (Bio-line) on ExicyclerTM 96 (Bioneer). qRT-PCR results were analyzed by Ct method for relative quantifications taking GAPDH or HPRT as internal controls.

### Immunofluorescent staining

The cells were fixed 4% paraformaldehyde (PFA; Sigma) for 30 min at room temperature and washed with phosphate-buffered saline (PBS) three times. Then, the cells were permeabilized with 0.2% TritonX-100 (Sigma) for 20 min at room temperature and washed with PBS three times. The cells were blocked in 3% BSA (Applichem) and 5% donkey serum (Merck) in PBS for 2 hours at RT. Immuno-labelling was performed with CDX2 (EPR2764Y, ABCAM) (1:250), KRT7 (M7018, DAKO) (1:100), OCT3/4 (sc-5279, SantaCruz) (1:100), NANOG (AB21624, ABCAM) (1:100), Mouse IgG (0.4 µg/mL) overnight at +4°C. Next day, the cells washed with 1X PBS for 3 times and they were incubated with appropriate secondary antibodies conjugated with Alexa-Flour 488, 555 or 568 for 3h at +4oC in dark. The cells were washed again with 1X PBS for 3 times. Images were acquired using a confocal microscope (Leica TCS SP8, USA) and processed via ImageJ.

### Immunoblotting

The cells were harvested in RIPA lysis buffer (150mM NaCl, 1% NP40, 0.5% sodium deoxycolate, 0.1% SDS, 50mM Tris pH 7,4) supplied with protease and phosphatase inhibitors (Roche, Switzerland). For the supernatants, TCA-acetone precipitation was performed. Protein samples were applied to SDS gel. Semidry transfer was performed by using Blotting papers (Sigma-Aldrich, USA) and PVDF membrane (Millipore, Ireland). After blocking with 5% non-fat dry milk, the membranes were incubated with primary antibodies (1:1000) +4°C overnight. Next day, the membrane was incubated with 1:2000 HRP-coupled secondary antibodies according to primary antibody’s host origin. Between all the steps, the membranes were washed with TBS-T for 3 times. Immunoblotted membranes were visualized with Syngene documentation system by using ECL HRP Substrate (Advansta, USA).

### Immunoassay

PGF was measured by Human PIGF ELISA (DPG00; RD Systems) according to manufacturer’s protocol. Diluted to appropriate concentration of samples and standards were transferred on 96 well-plate and incubated 2 hours at room temperature. The wells were washed four times with wash buffer and the conjugated antibody was added to the plate and incubated 2 hours at room temperature. Substrate Solution (1:1 Color Reagent A (H2O2): Color Reagent B (Tetramethylbenzidine)) was incubated 30 minutes at room temperature on dark. Finally, stop Solution (2M H2SO2) was added and mixed to stop the reaction and optical density was measured at 450 and 570 nm (VersaMax, Molecular Devices, USA).

### Plasmid construction and lentiviral transduction

The pCR-Blunt-II TOPO plasmid containing the NLRP7 cDNA was purchased from Imagenes. The region that encoded for NLRP7 cDNA was PCR amplified by using SalI and NotI restriction sites added primers and ligated into pENTR1A noc-cDB (W48-1). Next, NLRP7 and GFP genes were subcloned into pLEX307 (Addgene 41392) lentiviral expression vector using Gateway cloning technology (Invitrogen). Sequencing procedures were performed to verify no SNP or frameshift containing plasmids. NLRP7 or GFP expressing vectors were transfected to 293FT cells at a density of 2,5×106 in 10cm dishes to produce lentiviruses using Fugene (Promega). Virus containing medium were collected 2 days. PEG8000 (Sigma) was used to concentrate the virus titre. HM^iPSCs^ were infected with obtained viruses two times for 16h.

### RNA sequencing analysis

Total RNA was extracted using the Direct-zol RNA Isolation Kit (Zymogen) according to manufacturer’s protocol (n=2 in all cases). Library preparation using Truseq stranded mRNA LT Sample Prep Kit (Illumina) and RNA sequencing using Hiseq2500 (Illumina) was performed by Macrogen Inc. (Korea). RNA-seq data were processed and interpreted with the Genialis visual informatics platform (www.genialis.com). Briefly, RNA-seq reads were implemented to trimming (BBDuk), mapping (STAR), and expression quantification (featureCounts), respectively. Reads were mapped to Homo sapiens GRCh38 (Ensembl, version 92, ERCC). Differential gene expression analyses were performed using DESeq2 (Love, Huber and Anders, 2014). Lowly-expressed genes were filtered out from the differential expression analysis input matrix. The expression level (TPM, transcripts per kilobase per million) was determined by Cufflinks (http://cufflinks.cbcb.umd.edu). Heatmaps were built with Genialis platform according to row-wise scaled expression using Z-score based on TPM values of literature curated custom gene sets of general trophoblasts, CTs, EVTs or STs (Supplementary Table1, 2) (27– 29, 46, 47). Benporath-ES1 gene set were used as Pluripotency markers (48). The first 200 genes that are differentially upregulated in WTBAP cells relative to WTAP cells on day 2, are assigned as early BMP4 responsive genes (Supplementary Table3). Hierarchical clustering was performed using R, DEseq2 package to show sample distances. Violin plots were generated using Python 3.7.2 with Seaborn library. All of the RNA-seq data can be accessed with the GEO code GSE125592.

### Gene set enrichment analysis

Pre-ranked differentially expressed gene lists were implemented to GSEA using default settings (http://software.broadinstitute.org). For the identification of enrichments in pluripotency genes, BENPO-RATH ES1; placenta genes, Module38 and BMP pathway GO response to BMP modules were assessed.

### BMP4 Methylation Analysis

Genomic DNA was extracted using Quick-DNA − Miniprep Plus Kit (Zymo Research) according to manufacturer’s protocol. Bisulfite conversion of 500 ng DNA was conducted using EZ DNA Methylation-Gold− Kit (Zymo Research) as recommended by the manufacturer. Bisulfite modified DNA was amplified by PCR using HotStarTaq Master Mix (Qiagen). PCR and sequencing primers were designed by Pyromark Assay Design Software 2.0 (Qiagen) and available upon request. Biotinylated PCR products were mixed with streptavidin beads and converted to single strands using PyroMark Q96 Vacuum Workstation (Qiagen). Following annealing of sequencing primers, pyrosequencing was performed using Pyromark Q96 ID System (Qiagen). CpG methylation data were analyzed by PyroMark software. Samples were run in duplicate. Assays were verified using EpiTect unmethylated and methylated DNA (Qiagen).

### Statistics

Statistical analyses were performed by Graphpad Prism 6.0. (San Diago, USA). The results were implemented to Two-way ANOVA followed by Sidak’s multiple comparison test. Wilcoxon rank sum test was used to determine P values for violin plots. The bars are presented as the mean ±SD. P values of < 0.05 are considered statistically significant and represented as follows: *P ≤0.05, **P< 0.01 ***P < 0.005, ****P < 0.001. All the experiments were conducted at least three times unless indicated otherwise.

## ACKNOWLEDGEMENTS

We thank N.C.T. Emre (Bogazici University, Department of Molecular Biology and Genetics) for kindly providing pLEX307 plasmid. We are thankful to Onur Eskiocak for his help while creating most of the illustrations. We would like to thank Ahmet Kocabay and Ali Cihan Taskın for help with mouse experiments, Arzu Ruacan (Koç University, School of Medicine, Department of Pathology) for examination of histological sections. The authors gratefully acknowledge use of the services and facilities of the Koç University Research Center for Translational Medicine (KUTTAM), funded by the Republic of Turkey Ministry of Development. The content is solely the responsibility of the authors and does not necessarily represent the official views of the Ministry of Development. This work was supported by an EMBO Installation Grant (T.Ö.) and funded by The Scientific and Technological Research Council of Turkey (TUBITAK) (projects: SBAG-112S115 and KBAG-217z131) (N.Ö.), Bogazici University Research Funding Office (BAP:7360) (N.Ö.).

## Author Contributions

designed and carried out the experiments, analyzed the data. B.Ö. performed reprogramming experiments. A.A.G., N.Ö. and V.U. recruited the patient. I.S. helped with the qRT-PCR, BMP4 western blot and pyrosequencing experiments. A.A.G., T.T.Ö., N.Ö. designed the research and wrote the manuscript. T.T.Ö. and N.Ö. provided funding.

## Data deposition

RNA sequencing data have been deposited in the Gene Expression Omnibus (GEO) under accession code GSE125592.

