## Supplementary Data for "NLRP7 Plays A Functional Role in Regulating BMP4 Signaling During Differentiation of Patient-Derived Trophoblasts"

**This PDF file includes:**
Figs. S1 to S5
Captions for Databases S1 to S4

**Other supplementary materials for this manuscript include the following:**
Databases S1 to S4

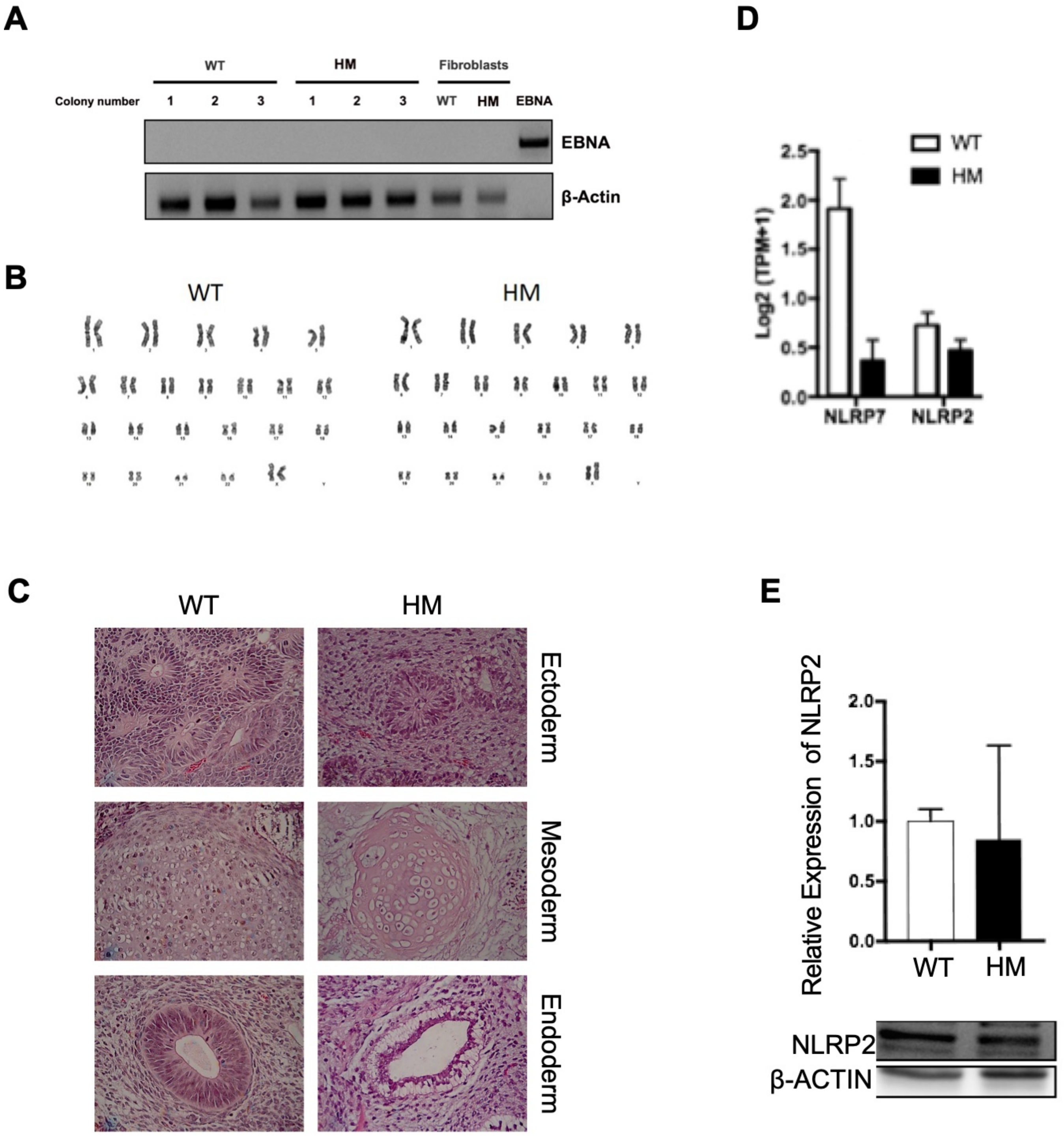


**Fig. S1.** (A) PCR-based EBNA integration test on genomic DNA isolated from three independent clones of each genotype. Plasmid was used as a positive control. (B) Karyotype of WT and HM iPSC lines showing normal chromosome complement. (C) Histological sections from in vivo teratoma formation assay showing that both iPSCs give rise to tissues of three embryonic germ layers; endoderm (glandular epithelium), ectoderm (neural rosettes), mesoderm (cartilage). (D) Log2 based expression values for NLRP7 and NLRP2 from RNA-sequencing; n=2 biological replicates. (E) NLRP2 mRNA and protein levels in $WT^{iPSC}$ and $HM^{iPSC}$ as assessed by RT-qPCR (top) and immunoblotting (bottom), n = 3 biological replicates. The bars represent mean ± SD. ***p< 0.001 by 2-way ANOVA followed by Sidak's multiple comparison test.

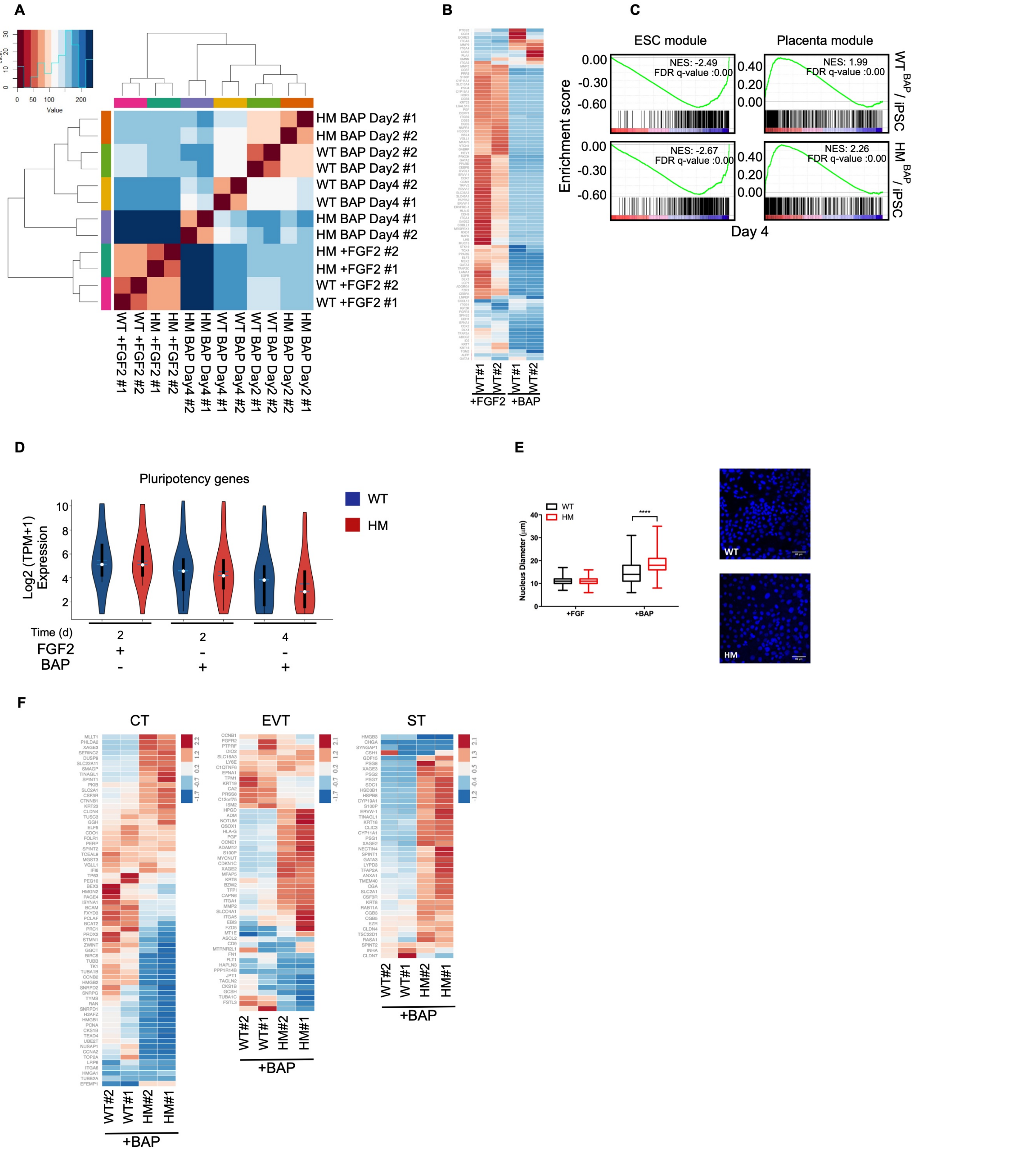


**Fig. S2.** (A) Hierarchical clustering depicting the transcriptome-wide sample distances. (B) Heatmap showing the expression of trophoblast markers across the time course of differentiation of WT (C) GSEA of the genes related with embryonic stem cells (ESC) or placenta. Genes were ranked according to log2 fold changes in gene expressions comparing BAP treated cells to FGF2 treated control cells on day 4. (D) Violin plot of pluripotency genes (n=2, p value <0.05, Wilcoxon rank sum test). (E) Nucleus sizes of DAPI stained cells were measured by ImageJ. n=100. Scale bar;10μM., ***p < 0.005 by 2way ANOVA followed by Sidak's multiple comparison test. (F) Heatmaps of genes for trophoblast sub-types; cytotrophoblast (CT), extravillous trophoblast (EVT), syncytiotrophoblast (ST).

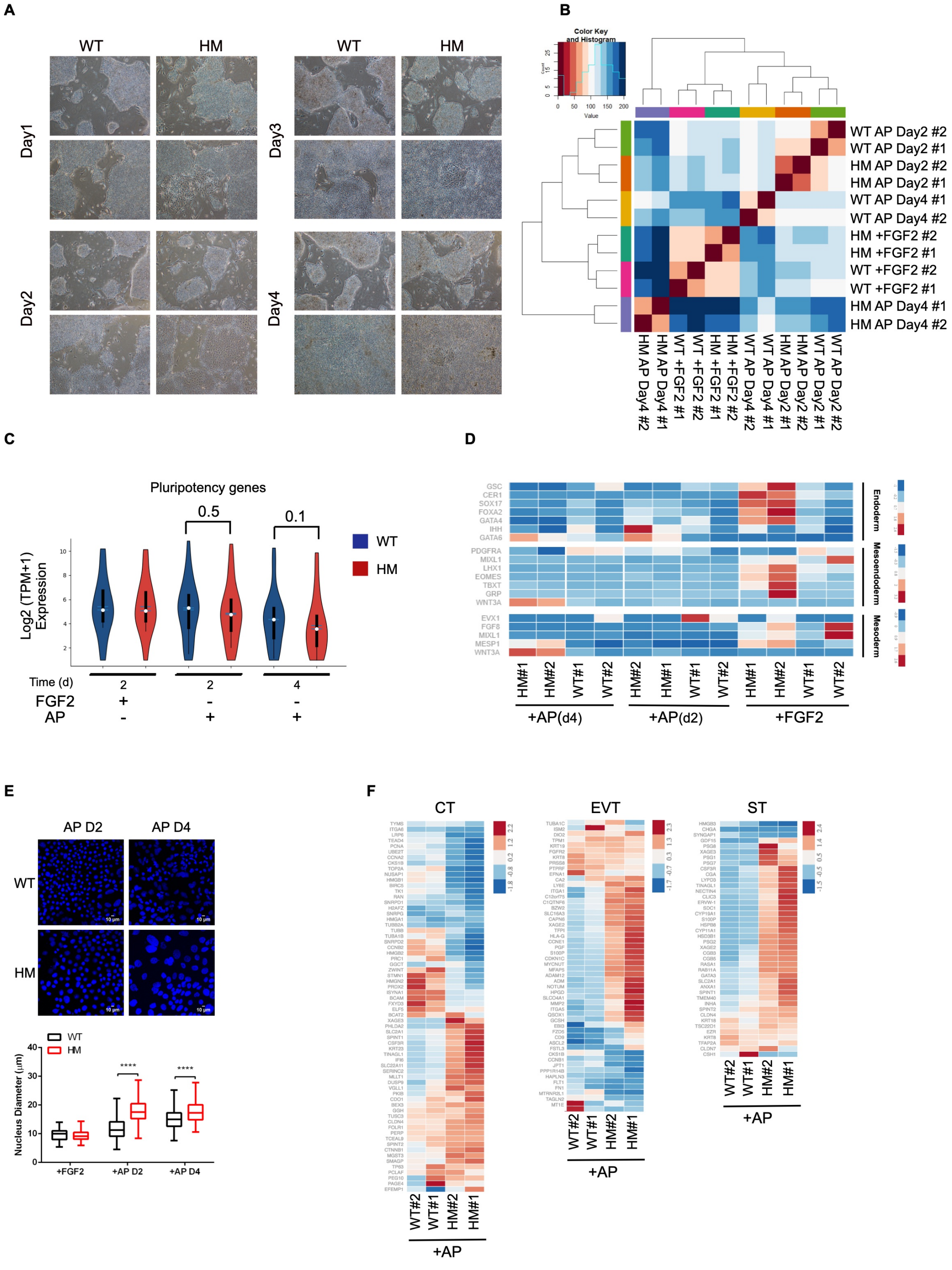

**Fig. S3.** (A) Changes in colony morphologies upon AP exposure were shown by light microscopy with 4x magnification. (B) Hierarchical clustering depicting the transcriptome-wide sample distances. (C) Violin plot of pluripotency genes (n=2, p <0.05, Wilcoxon rank sum test).(D) Heatmaps showing the expression (z-score) of endoderm , mesoendoderm and mesoderm markers. (E) Nucleus sizes of DAPI stained cells were measured by ImageJ. n=100, Scale bar; 10µM. ***p < 0.005 by 2way ANOVA followed by Sidak's multiple comparison test. (F) Heatmaps showing the expression (z-score) of trophoblast sub-types; cytotrophoblast (CT), extravillous trophoblast (EVT), syncytiotrophoblast (ST).

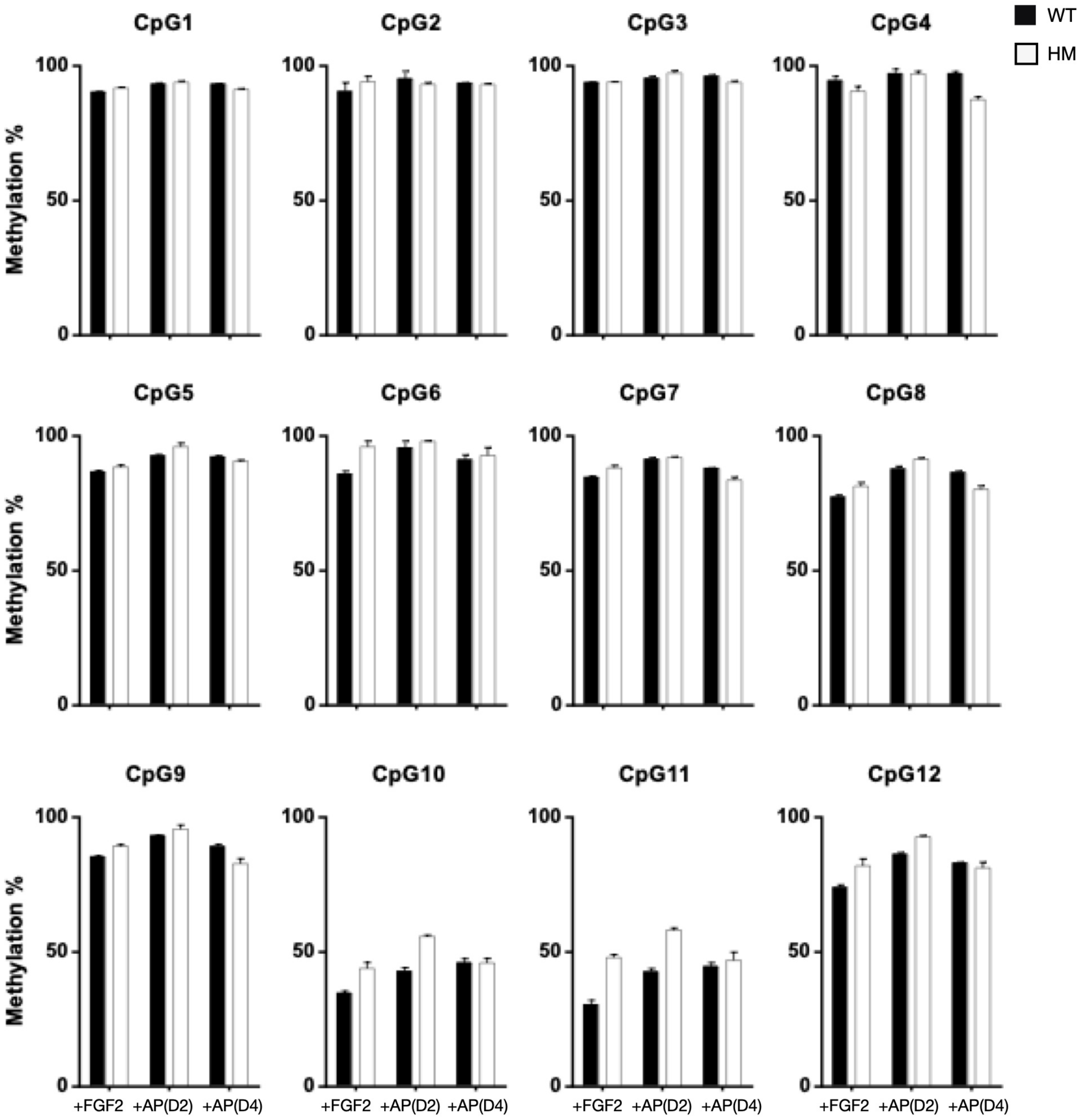


**Fig. S4.** Methylation status of BMP4 during time course of trophoblast differentiation. The percentage of methylation at 12 different CpG sites in the promoter and gene body of BMP4 were assayed by pyrosequencing upon AP exposure. The bars represent mean percentage of methylation, ± SD, n = 2 biological replicates.

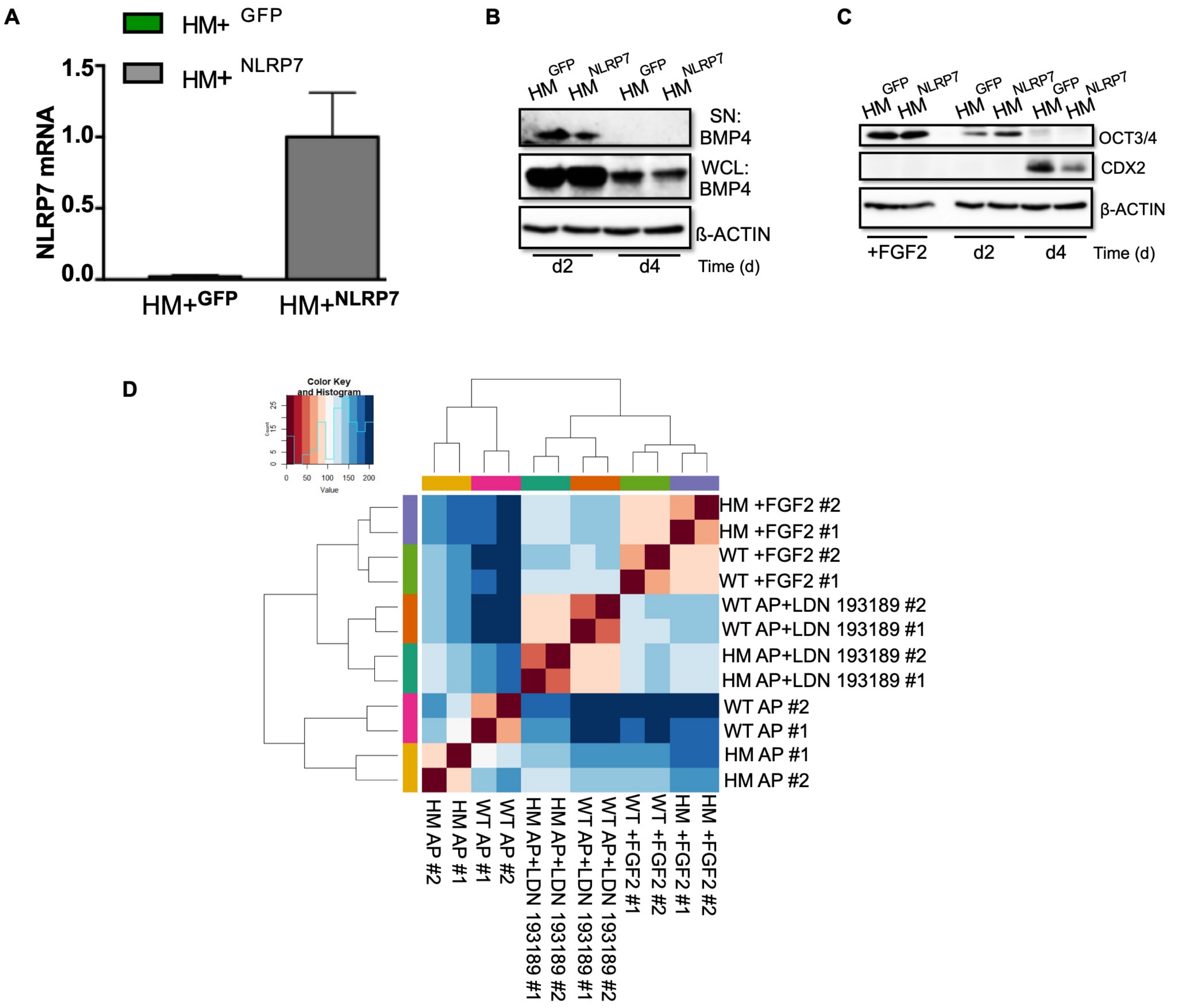


**Fig. S5.** (A) NLRP7 overexpression in HM cells by lentiviral transduction. RT-qPCR showing NLRP7 over-expression. GFP expressing cells were used as a control. Relative mRNA levels were normalized to GAPDH. The bars represent mean fold change (2△△ CT) ± SD, n = 3 biological replicates. (B) Western blotting of BMP4. Supernatant, SN; whole cell lysate, WCL. (C) Immunobloting for CDX2, OCT3/4. $\beta$-actin was used as a loading control. (D) Hierarchical clustering showing the transcriptome-wide sample distances.

**Additional data table S1 (ST1.xlsx)**

List of general trophoblast markers. The compiled list from recent literature for general trophoblast markers is provided.

**Additional data table S2 (ST2.xlsx)**

List of trophoblast sub-type markers: cytotrophoblasts (CT), extravillous trophoblasts (EVT) and syncytiotrophoblasts (ST). Trophoblast sub-type markers are listed according to recent literature.

**Additional data table S3 (ST3.xlsx)**

List of early BMP4-responsive genes. The first 200 number of genes that are upregulated in the transcriptomes of BAP-treatment in comparison to AP-treatment of WT cells on day 2 are listed.

**Additional data table S4 (ST4.xlsx)**

List of qRT-PCR primers. The sequences of primers used in gene expression studies are listed.
